# Epigenetic changes induced by *Bacteroides fragilis* toxin (BFT)

**DOI:** 10.1101/301374

**Authors:** Jawara Allen, Stephanie Hao, Cynthia L. Sears, Winston Timp

## Abstract

Enterotoxigenic *Bacteroides fragilis* (ETBF) is a gram negative, obligate anaerobe member of the gut microbial community in up to 40% of healthy individuals. This bacterium is found more frequently in people with colorectal cancer (CRC) and causes tumor formation in the distal colon of mice heterozygous for the <u>a</u>denomatous <u>p</u>olyposis <u>c</u>oli gene (*Apc*^+/−^); tumor formation is dependent on ETBF-secreted *Bacteroides fragilis* toxin (BFT). Though some of the immediate downstream effects of BFT on colon epithelial cells (CECs) are known, we still do not understand how this potent exotoxin causes changes in CECs that lead to tumor formation and growth. Because of the extensive data connecting alterations in the epigenome with tumor formation, initial experiments attempting to connect BFT-induced tumor formation with methylation in CECs have been performed, but the effect of BFT on other epigenetic processes, such as chromatin structure, remains unexplored. Here, the changes in chromatin accessibility (ATAC-seq) and gene expression (RNA-seq) induced by treatment of HT29/C1 cells with BFT for 24 and 48 hours is examined. Our data show that several genes are differentially expressed after BFT treatment and these changes correlate with changes in chromatin accessibility. Also, sites of increased chromatin accessibility are associated with a lower frequency of common single nucleotide variants (SNVs) in CRC and with a higher frequency of common differentially methylated regions (DMRs) in CRC. These data provide insight into the mechanisms by which BFT induces tumor formation. Further understanding of how BFT impacts nuclear structure and function in vivo is needed.

**Importance:** Colorectal cancer (CRC) is a major public health concern; there were approximately 135,430 new cases in 2017, and CRC is the second leading cause of cancer-related deaths for both men and women in the US (1). Many factors have been linked to CRC development, the most recent of which is the gut microbiome. Pre-clinical models support that enterotoxigenic *Bacteroides fragilis* (ETBF), among other bacteria, induce colon carcinogenesis. However, it remains unclear if the virulence determinants of any pro-carcinogenic colon bacterium induce DNA mutations or changes that initiate clonal CEC expansion. Using a reductionist model, we demonstrate that BFT rapidly alters chromatin structure and function consistent with capacity to contribute to CRC pathogenesis.

## Introduction

*Bacteroides fragilis* is a gram-negative, obligate anaerobe that is found consistently, but in low numbers, in the gut microbial community (2). ETBF is a particular subtype of *Bacteroides fragilis*, characterized by its production of *Bacteroides fragilis* toxin (BFT) and its association with diarrhea, inflammatory bowel disease and colon cancer(3, 4). Since the identification of ETBF, many studies have examined the carriage rate of this potentially pathogenic bacterium. Some studies have probed the mucosa for the bacterium and found an asymptomatic carriage rate as high as 67% (4), but most studies examine the stool and report rates of 10%-12% (3). So, while we do not yet know exactly how many people are colonized with ETBF, it is clear that a significant fraction of the population likely experiences prolonged exposure to the bacterium, and thus BFT.

The *bft* gene codes for the BFT pre-pro-protein that is processed by ETBF, yielding a secreted active 20 kD zinc-dependent metalloprotease (3). There are three isotypes of the BFT protein, each produced by a different *bft* gene (3), with the most potent of the BFT isotypes being BFT2. Expression of BFT2 is required for ETBF-induced tumorigenesis in multiple intestinal neoplasia (*Apc^min/+^*) mice (5). When interacting with colon epithelial cells (CECs), BFT2 binds to an unidentified cell surface receptor which leads to rapid, but indirect, cleavage of E-cadherin that, in turn enhances barrier permeability *in vivo* (6). Because a pool of β–catenin is associated with the intracellular domain of E-cadherin, cleavage of E-cadherin triggers release of β–catenin and its subsequent nuclear localization with upregulation of c-Myc expression and enhanced cellular proliferation (7).

Though we know some of the immediate downstream effects of BFT on CECs, we still do not understand how this potent exotoxin causes lasting changes in CECs that lead to tumor formation and growth. A possible explanation is an epigenetic progenitor model to cancer, in which epigenetic dysregulation enables tumor cell survival (8). Epigenetics is the study of heritable phenotypic variation that is not caused by base pair changes in the genome. Epigenetic changes have been linked to various cancers; in colorectal cancer specifically, hypermethylation of CpG islands and global hypomethylation have both been implicated in tumor development and progression (9) and global dysregulation of the methylome has been observed in clinical colon cancer samples (10).

Initial experiments exploring the impact of BFT on the epigenome have been promising. By tracking the localization of specific epigenetic regulators, one study showed that inoculation of C57BL/6J mice with ETBF causes an upregulation of gene silencing complexes at CpG islands (11). Subsequent studies have shown that inoculation of Apc^*min/+*^ mice with ETBF also causes recruitment of DNA methyltransferase 1 (DNMT1), a recruitment potentially mediated by DNA mismatch repair proteins (12). While these studies have established a role for gene regulation and DNA methylation in the effect of ETBF on CECs, studies do not report on how ETBF affects other epigenetic processes, such as chromatin structure. This is of particular importance because chromatin state as measured by changes in various histone marks has been linked to colon cancer development (9), and differences in chromatin structure have been shown to impact mutation rates along the genome (13)(14).

Herein, we further test the hypothesis that the epigenome is at least one mechanism by which BFT acts to increase colon tumor formation. To test our hypothesis, we use a reductionist model system to investigate the effect of BFT2 on CEC chromatin structure. Specifically, we examined whether BFT2 alters chromatin accessibility and subsequently, gene expression, using a model human colon epithelial cell line, HT29/C1, that is known to be uniformly sensitive to BFT2 and for which extensive prior data exist (3). We extended our analyses to seek to determine if BFT2-induced chromatin changes were associated with functional consequences by correlation with transcription factor binding motifs and known cancer-causing mutations in human CRC.

## Results

### BFT2 induces dynamic changes in chromatin accessibility throughout the genome

We initially examined time points of 24 and 48 hours after BFT treatment (fig 1A), performed in biological triplicate. To assay chromatin state, we performed ATAC-seq (<u>A</u>ssay for <u>T</u>ransposase-<u>A</u>ccessible <u>C</u>hromatin using <u>seq</u>uencing) on BFT2-treated and untreated cells at each time point (15). Briefly, ATAC-seq leverages the accessibility of chromatin to preferentially “attack” open chromatin using a transposase (Nextera; Illumina), resulting in a tagmentation reaction. Subsequent preferential amplification and downstream sequencing results in a distribution of sequencing reads which reveals areas of open versus closed chromatin by examination of the depth of coverage (i.e., the number of reads which align to each area of the genome). We utilized software tools like MACS2 (see Methods) to detect peaks in the coverage that indicate open chromatin regions.

**Figure 1:**
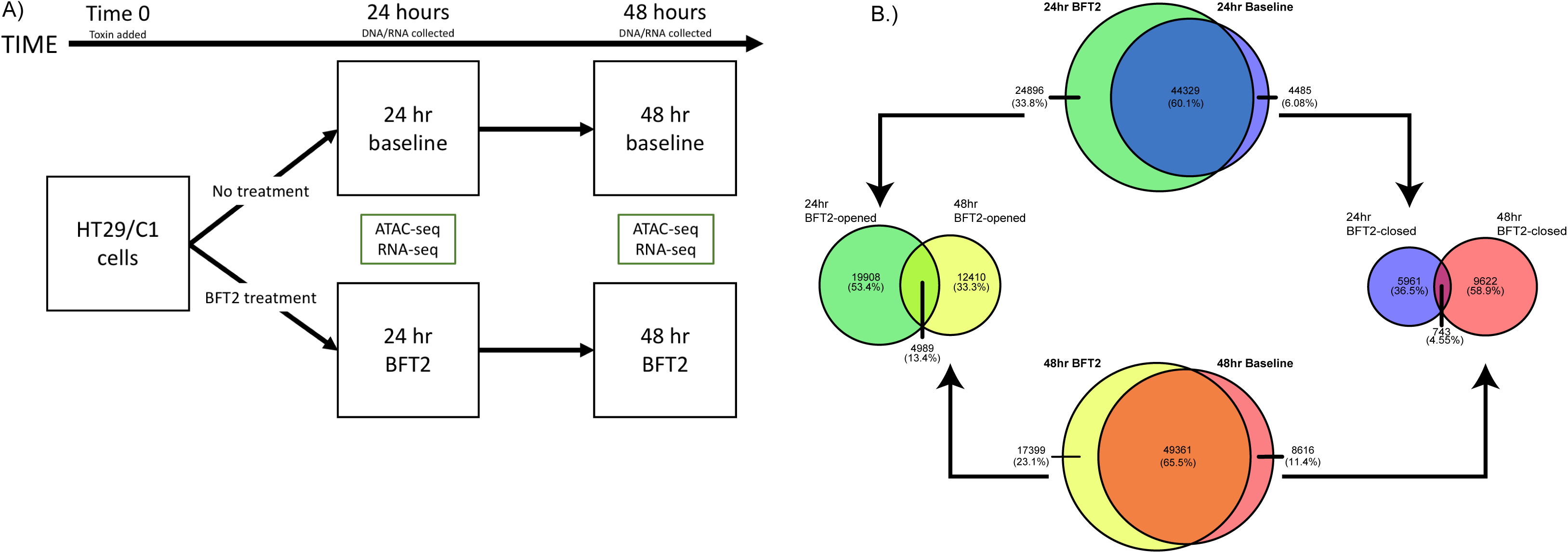
BFT2 induces dynamic changes in chromatin accessibility. A.) HT29/C1 cells were treated with BFT2 or left untreated for 24 or 48 hours. After each time point, DNA and RNA were isolated. DNA was used to perform ATAC-seq, a measure of genome-wide chromatin accessibility. RNA was used to perform RNA-seq, a measure of genome-wide gene expression. B.) Venn diagrams showing the overlap of areas of accessible chromatin in untreated cells (baseline) and BFT2-treated cells at 24 hours (top) and 48 hours (bottom). Areas of accessible chromatin detected after BFT2 treatment at 24 or 48 hours, but not in untreated cells, were categorized as “BFT2-opened” while areas of accessible chromatin detected in untreated cells, but not after BFT2 treatment, were categorized as “BFT2-closed.” Left and right Venn diagrams show the overlap of BFT2-opened (left) or BFT2-closed (right) regions of the genome at 24 and 48 hours. For all Venn diagrams, the number of open peaks (chromatin accessible) detected and the percentage of those peaks in each sector of the Venn diagrams are shown. Also, labels (bold) at the top of each Venn diagram describe the entire circle, both overlapping and nonoverlapping regions, for each experimental condition.

We first quantified the total number of peaks of chromatin accessibility in our baseline samples (untreated cells) and BFT2-treated samples. We then classified these peaks into 2 types at 24 and 48 hours: BFT2-opened peaks (peaks of chromatin accessibility detected only after BFT2 treatment), and BFT2-closed peaks (peaks of chromatin accessibility that disappeared after BFT2 treatment). This initial analysis revealed that only a fraction (40% at 24 hours and 35% at 48 hours) of the peaks identified by ATAC-seq are affected by BFT2 treatment, and that BFT2 treatment appears more likely to open chromatin than to close it (~11:3 ratio at 24 hours; ~5:3 ratio at 48 hours) (fig 1B). We next analyzed the overlap of our BFT2-opened peaks and BFT2-closed peaks between 24 and 48 hours in order to determine if BFT2-induced changes in chromatin accessibility are stable. We found that only 13.4% (4989 peaks) of 24 hours and 48 hours BFT2-opened peaks overlap, and 4.55% (743 peaks) of 24 hours and 48 hours BFT2-closed peaks overlap (fig 1B). These results emphasize the dynamic nature of BFT2-induced changes of chromatin accessibility.

To explore how BFT2 affects peak distribution throughout the genome, we performed a detailed analysis on the BFT2-opened and BFT2-closed chromatin regions (i.e. the regions of the genome affected by BFT2 treatment). When we analyze the distribution of these peaks, we see that more of BFT2-opened and BFT2-closed peaks are located within intergenic regions and coding regions than promoter regions (Table 1). To determine if this distribution was a byproduct of the ATAC-seq assay or a specific effect of BFT2, we compared those peak distributions to the distribution present in our baseline and BFT2-treated samples. This analysis reinforced that more of the BFT2-opened and BFT2-closed peaks are located within intergenic and coding regions, while less are located within promoter regions (Table 1). Although we expect that the effect in all three regions is important, these results suggest that chromatin accessibility is affected differentially across the genome by BFT2.

**Table 1:** Distribution of ATAC-seq peaks throughout the genome.

|  | 24hr<br>Baseline<br>(48,982)<sup>a</sup> | 24hr<br>BFT2<br>(69,415) | 48hr<br>Baseline<br>(58,147) | 48hr<br>BFT2<br>(66,945) | 24hr<br>BFT2-Opened<br>(24,967) | 24hr<br>BFT2-Closed<br>(6,754) | 48hr<br>BFT2-Opened<br>(17,459) | 48hr<br>BFT2-Closed<br>(10,417) |
| --- | --- | --- | --- | --- | --- | --- | --- | --- |
| **Promoter**<br>**(0.5%)<sup>b</sup>** | 41.6% | 33.7% | 37.9% | 34.9% | 17.4% | 18.2% | 16.0% | 16.7% |
| **Coding**<br>**(44.3%)** | 26.8% | 31.5% | 28.7% | 30.7% | 40.2% | 37.4% | 41.1% | 39.4% |
| **Intergenic**<br>**(55.0%)** | 31.6% | 34.8% | 33.4% | 34.4% | 42.4% | 44.4% | 42.9% | 43.9% |

The coding region includes 5' UTR, introns and exons
<sup>a</sup> Number of peaks per sample in parentheses
<sup>b</sup> Percent of the human genome represented by each region in parentheses

### BFT2 treatment induces changes in gene expression that are more pronounced at 24 hours than 48 hours

To examine the relationship between gene expression and chromatin state, we performed RNA-seq on the same samples used for ATAC-seq analysis. While the effect of BFT2 treatment on select genes at relatively early time points has been studied (16, 17), how this toxin affects genome wide expression and whether these effects persist is still unknown. After treatment with BFT2 for 24 hours, 70 genes were differentially expressed (P-value < 0.01). Of these genes, 41 showed a decrease in gene expression, while 29 showed an increase in gene expression (fig 2A, supplementary table 1). After BFT2 treatment for 48 hours, we found only 16 differentially expressed genes (P-value < 0.01); 3 showed a decrease in gene expression and 13 showed an increase in gene expression (fig 2B, supplementary table 2). This result follows a trend seen throughout our analyses and the literature (7, 16, 18) in which BFT2 has a more potent effect on CECs at 24 hours after toxin treatment that seems to diminish by 48 hours. A lineplot of the top 16 differentially expressed genes at 24 and 48 hours shows that gene expression at the two time points tends to differ; genes differentially expressed at 24 hours tend to move closer to baseline levels by 48 hours, while genes differentially expressed at 48 hours tend to have expression levels closer to baseline at 24 hours (supplementary fig 1). Notably, none of the genes differentially expressed at 24 hours show statistically significant differential expression at 48 hours as well. These data reinforce the observation that BFT2-induced gene expression changes are highly dynamic, and time specific, similar to BFT2-induced changes in chromatin accessibility.

**Figure 2:**
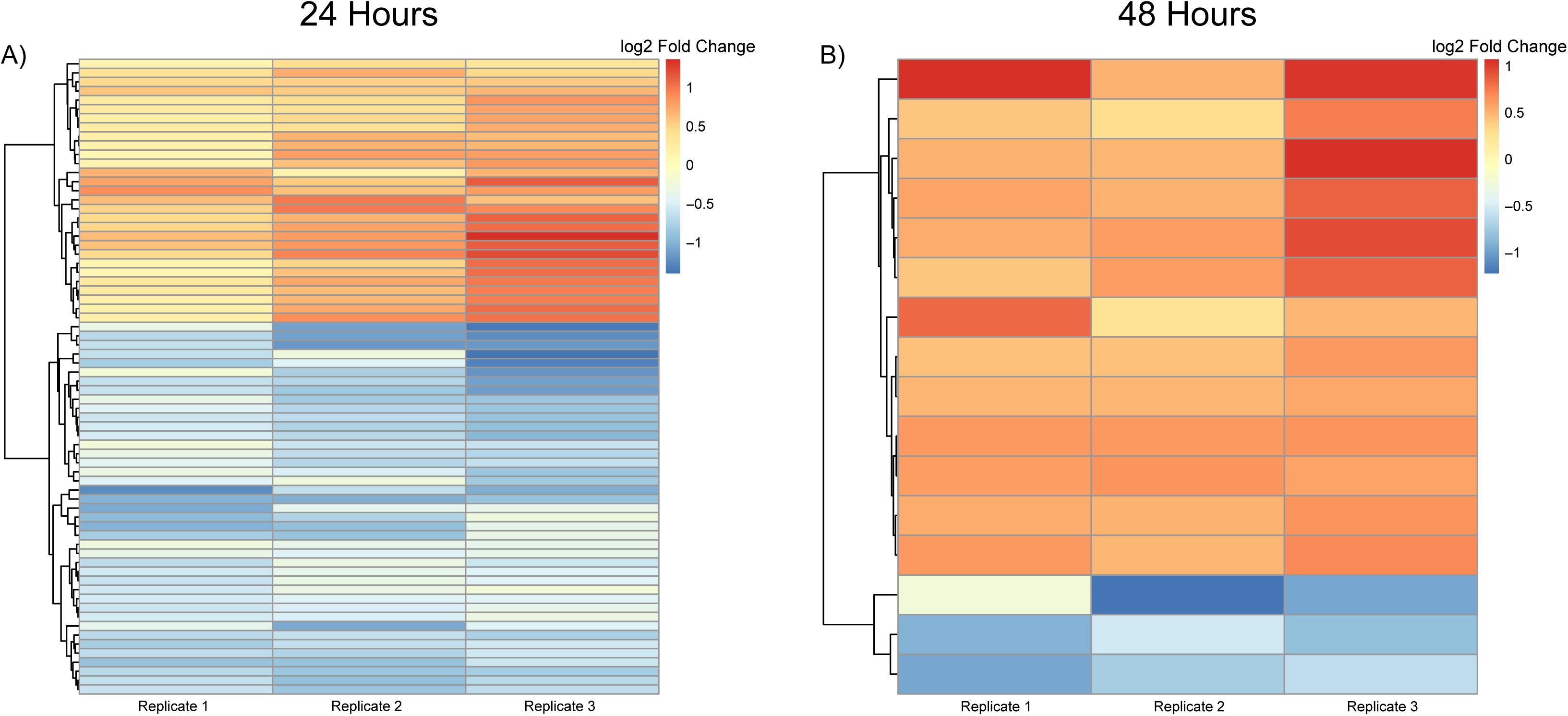
The effects of BFT2 on gene expression are more pronounced 24 hours after treatment than 48 hours. RNA-seq was performed on untreated cells and those treated with BFT2 in order to determine if BFT2 alters gene expression in HT29/C1 cells. A) Heatmap showing the change in gene expression at 24 hours after BFT2 treatment for those genes with P-values < 0.01. B) Heatmap showing the changes in gene expression at 48 hours after BFT2 treatment for those genes with P-values < 0.01. For both heatmaps, colors represent Log2FC (fold change) values of gene expression in BFT2-treated samples compared to gene expression in baseline samples. Full list of differentially expressed genes after BFT2 treatment are in supplementary tables 1 and 2. C.) Dotplot showing the output from a gene set enrichment analysis looking for gene ontology (GO) and reactome gene sets that are upregulated or downregulated at 24 or 48 hours after BFT2 treatment. Only gene sets with an FDR qval < 0.25 are shown.

To perform a Gene Set Enrichment (GSE) analysis on the list of genes that were differentially expressed at 24 hours and 48 hours after BFT2 treatment, the cutoff for significance was relaxed to a P-value of less than 0.05. The GSE analysis revealed several processes that were overrepresented in our list of upregulated and downregulated genes (fig 2c). Specifically, at 24 hours after BFT2 treatment, expression of transcription factors involved in nucleic acid binding were downregulated. At 48 hours after BFT2 treatment, genes related to the cell cycle, chromosome organization and response to DNA damage, among others, were upregulated (fig 2C).

### Chromatin accessibility correlates with gene expression at baseline and at 24 hours after BFT2 treatment

We next tested the hypothesis that BFT2-induced changes in chromatin accessibility correlate with BFT2-induced changes in gene expression. First, we classified genes by promoter chromatin accessibility status in our 24hr untreated (baseline) sample. Only promoter peaks were used because of the intuitive connection between open promoter chromatin and changes in gene expression. Those genes that contained an ATAC-seq peak overlapping their promoter region were categorized as “open” and those without a peak were categorized as “closed”. We then extracted normalized gene expression values (transcripts per million-TPM) for each of these genes to determine if the average gene expression values were different for the two groups. We found that on average, gene expression was statistically higher among genes that were “open” in our 24hr baseline sample compared to those that were “closed” (P-value < 2e^−16^) (fig 3A).

**Figure 3:**
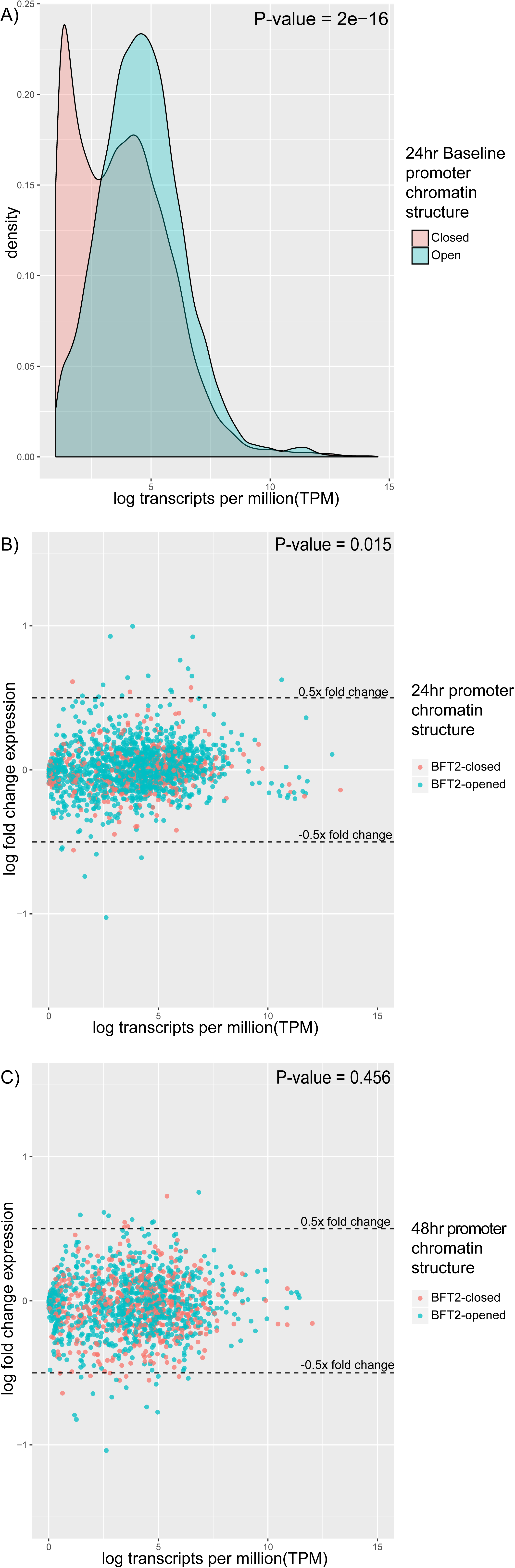
Chromatin accessibility and gene expression correlate at baseline and 24 hours after BFT2 treatment. A.) Stacked histogram showing the correlation between gene expression and chromatin accessibility at baseline. Genes were first sorted by the presence of a peak in their promoter in the 24hr baseline sample. Genes with a promoter peak were considered “open” while those without a peak were considered “closed.” Log transcripts per million (TPM) gene expression values were then extracted and compared. The P-value was calculated by comparing the gene expression of genes with a promoter peak to those without a promoter peak in the 24hr baseline sample. B-C) MA plot analyzing gene expression change after BFT2 treatment at 24 and 48 hours. Genes are colored according to the presence of a BFT2-opened peak or a BFT2-closed peak in their promoter region. The dotted lines represent a log2 fold change in expression of +/− O.5. The P-values were calculated by comparing the fold change in gene expression for genes with a promoter BFT2-opened peak to those with a promoter BFT2-closed peak. For all graphs, P-values were calculated using the Wilcoxon rank sum test.

Next, we examined the correlation between changes in chromatin accessibility and changes in gene expression at the same time points. To do this, we again focused on BFT-opened and BFT-closed peaks which overlapped with gene promoter regions. We then calculated the logarithmic fold-change (Log2FC) expression values for each of these genes at 24 or 48 hours after BFT2 treatment. We found that genes with a BFT2-opened peak overlapping their promoter had a three-fold higher Log2FC in gene expression after 24 hours of BFT2 treatment than genes with a BFT2-closed peak at the same time point. At 48 hours after BFT2 treatment, we found no difference in Log2FC gene expression (fig 3 B-C). A similar analysis can be performed for the 86 genes that show statistically significant differences in gene expression after BFT2 treatment (fig 2 A-B, Supplementary tables 1, 2). Among those, 7 contained BFT2-opened peaks overlapping their promoter (6 at 24 hours and 1 at 48 hours). Of these genes, 5 showed an increase in gene expression, while 2 showed a decrease. No BFT2-closed peaks overlapped with the promoters of these 86 genes at either time point (supplementary tables 1, 2). Taken together, these results suggest that at 24 hours after BFT2 treatment, changes in chromatin structure largely correlate with changes in gene expression, but this relationship disappears by 48 hours.

### Treatment with BFT2 causes increased chromatin accessibility at transcription factor binding sites

Gene expression is regulated via multiple mechanisms, including the binding of transcription factors to enhancers and gene promoters. Therefore, we also wanted to test the hypothesis that BFT2-induced changes in chromatin accessibility impact transcription factor binding sites and thus gene expression. This can be done by looking for specific transcription factor motifs at sites of altered chromatin accessibility, or by calculating the overlap of sites of altered chromatin accessibility with sites of transcription factor binding. This latter approach is only available for a handful of transcription factors, such as CTCF, which have been extensively profiled via Chip-seq experiments in various cell types (19).

We first used the haystack_bio bioconda package to query our ATAC-seq data for specific transcription factor motifs. This analysis revealed several transcription factor motifs that are enriched in our BFT2-treated or baseline samples (Tables 2,3; Supplementary tables 5,6). In contrast to the BFT2-dependent dynamic chromatin changes identified in our earlier analyses (fig 1B, Table 1), many of the most enriched transcription factor motifs identified are present after both 24 and 48 hours of BFT2 treatment. Notably, these include JUND, JDP2, FOSL1, JUNB, FOS, FOS:JUN and ZBTB33, many of which are downstream of mitogen-activated protein kinase (MAPK) pathways previously shown to be modulated by BFT2 (18, 20, 21). In subsequent analyses, we specifically examined FOSL1, as it was the only transcription factor whose expression was upregulated 24 hours after BFT2 treatment. We noted that while several FOSL1-regulated genes were also upregulated after BFT2 treatment, a near equal proportion were also downregulated (supplementary tables 3,4). For FOSL1 to augment gene expression, it needs to bind to an accessible DNA-binding site, so we next examined the FOSL1-regulated genes and classified them according to accessibility 24 hours after BFT2 treatment, as determined by ATAC-seq. We found that 6 of 7 genes with accessible DNA binding sites showed an increase in expression after BFT2 treatment, and only 1 of 6 genes without an accessible DNA binding site showed an increase in expression after BFT2 treatment (supplementary tables 3,4).

**Table 2:** Transcription factor binding motifs enriched after cell treatment with BFT2 for 24 hours.

| Motif Name | Presence in target | Presence in background | Ratio | P-value |
| --- | --- | --- | --- | --- |
| **JUNB** | 12.68% | 10.17% | 1.22 | 1.90E-40 |
| **FOS** | 12.40% | 9.97% | 1.22 | 8.14E-39 |
| **JUN(var.2)** | 12.89% | 10.43% | 1.22 | 1.71E-38 |
| **JUND** | 11.97% | 9.59% | 1.22 | 2.45E-38 |
| **FOSL1** | 12.50% | 10.09% | 1.22 | 1.27E-37 |
| **FOS::JUN** | 12.11% | 9.75% | 1.22 | 2.74E-37 |
| **JDP2** | 10.17% | 8.01% | 1.24 | 8.26E-37 |
| **BATF::JUN** | 10.91% | 8.82% | 1.21 | 2.48E-32 |
| **NFE2** | 10.47% | 8.43% | 1.22 | 7.61E-32 |
| **ZBTB33** | 12.90% | 15.27% | 0.85 | 4.76E-31 |
| **<sup>a</sup>CTCF** | 37.76% | 36.92% | 1.02 | 3.38E-03 |

Only top 10 motifs are shown. Full list in supplementary table 5
P-values calculated using Fisher's exact test
Ratio < 1.0 represents enrichment in baseline sample
<sup>a</sup>included due to physiological relevance

**Table 3:** Transcription factor binding motifs enriched after cell treatment with BFT2 for 48 hours.

| Motif Name | Presence in target | Presence in background | Ratio | P-value |
| --- | --- | --- | --- | --- |
| **ZBTB33** | 13.59% | 14.79% | 0.92 | 1.84E-09 |
| **FOS** | 11.26% | 10.26% | 1.09 | 1.49E-08 |
| **JUNB** | 11.43% | 10.43% | 1.09 | 1.51E-08 |
| **JUND** | 10.78% | 9.82% | 1.09 | 2.32E-08 |
| **FOS::JUN** | 10.95% | 10.01% | 1.09 | 7.19E-08 |
| **JDP2** | 9.20% | 8.34% | 1.09 | 7.80E-08 |
| **FOSL1** | 11.27% | 10.35% | 1.08 | 1.60E-07 |
| **E2F4** | 21.39% | 22.60% | 0.95 | 2.60E-07 |
| **JUN(var.2)** | 11.70% | 10.82% | 1.07 | 1.06E-06 |
| **NFE2** | 9.60% | 8.80% | 1.08 | 1.21E-06 |

Only top 10 motifs are shown. Full list in supplementary table 5
P-values calculated using Fisher's exact test
Ratio < 1.0 represents enrichment in baseline sample

While changes in promoter chromatin structure have been most extensively investigated, and provide many avenues for exploration, our data indicate that BFT2-induced changes in chromatin structure occur more frequently in intergenic regions. We hypothesized that these intergenic chromatin accessibility changes may affect chromatin architecture more globally and sites of CTCF binding specifically. CTCF is a DNA binding protein that regulates global chromatin structure by mediating chromatin looping via binding to and bringing together distant regions of the genome (19). The CTCF motif was enriched in our ATAC-seq data after 24 hours, but not 48 hours, of BFT2 treatment (Tables 2, 3). To further investigate this enrichment, we calculated a jaccard index for the overlap of CTCF binding sites in CACO-2 cells and HT29/C1 cells (the latter using our ATAC-seq data); both cell lines are human transformed colon epithelial cell lines. We found that after 24 hours of BFT2 treatment, but not 48 hours, the HT29/C1/CACO-2 jaccard index was increased by 1.13 fold (data not shown). These results again suggest that sites in the genome that show increases in chromatin accessibility after BFT2 treatment are more likely to also be sites of CTCF binding.

### Chromatin accessibility is associated with differential DNA methylation and DNA mutation

The previous analyses help us better understand how BFT2 alters chromatin accessibility in CECs and to connect chromatin changes with anticipated gene expression changes. However, they do not help us understand how changes in chromatin accessibility may contribute to common DNA-modifications found in CRC. To explore this, we looked for a correlation between BFT2-induced changes in chromatin accessibility and single nucleotide variants (SNVs) and common sites of differential methylation (DMRs) in CRC. SNVs and DMRs were extracted from the COSMIC database. We calculated the proportion of peaks in each sample that overlapped with a common SNV or DMR. We then used a chi-square test to compare our treated and untreated samples in order to determine if the proportion of peaks overlapping a SNV or DMR differed after treatment of HT29/C1 cells with BFT2.

Correlating our chromatin accessibility changes with SNVs in CRC allows us to explore the effects of BFT2-induced chromatin accessibility changes in coding regions of the genome, where specific chromatin structure states have been linked to decreased mutation rates in cancer (13). In contrast, examining DMRs allows us to explore whether BFT2-induced chromatin accessibility is associated with hypermethylation of CpG islands in promoter regions, or hypomethylation of short interspersed nucleotide elements (SINEs) or long interspersed nucleotide elements (LINEs) in intergenic regions.

Our SNV analysis revealed a smaller proportion of peaks overlapping with common SNVs in CRC 24 hours [OR(95%CI): 0.819(0.794-0.844)] and 48 hours [OR(95%CI): 0.964(0.936-0.993)] after BFT2 treatment (Table 4). Similar to our prior data, the effect of BFT2 was more pronounced 24 hours after cell treatment. Thus, contrary to our hypothesis, these data suggest that regions of the genome with BFT2-induced increases in chromatin accessibility are *less* likely to contain SNVs commonly found in CRC.

**Table 4:** Association between BFT2-induced chromatin accessibility changes and common Single Nucleotide Variants (SNVs)/Differentially Methylated Regions (DMRs) in CRC.

|  | Number of overlapping peaks | Number of non-overlapping peaks | Total number of peaks | Odds<sup>a</sup> | Odds ratio<sup>b</sup> (95% CI<sup>c</sup>) |
| --- | --- | --- | --- | --- | --- |
| **SNVs** |  |  |  |  |  |
| **24hr BFT2** | 11063 | 58352 | 69415 | .0281 | 0.818 (0.794-0.844) |
| **24hr Baseline** | 9210 | 39772 | 48982 | .0271 |  |
| **48hr BFT2** | 11147 | 55798 | 66945 | .0259 | 0.964 (0.936-0.993) |
| **48hr Baseline** | 9981 | 48166 | 58147 | .0237 |  |
| **DMRs** |  |  |  |  |  |
| **24hr BFT2** | 1900 | 67515 | 69415 | .0281 | 1.037 (0.965-1.114) |
| **24hr Baseline** | 1294 | 47688 | 48982 | .0271 |  |
| **48hr BFT2** | 1692 | 65253 | 66945 | .0259 | 1.094 (1.018-1.177) |
| **48hr Baseline** | 1346 | 56801 | 58147 | .0237 |  |
| **DMRs hypermethylated** |  |  |  |  |  |
| **24hr BFT2** | 944 | 68471 | 69415 | .0138 | 0.979 (0.887-1.082) |
| **24hr Baseline** | 680 | 48302 | 48982 | .0141 |  |
| **48hr BFT2** | 953 | 65992 | 66945 | .0144 | 1.101 (1.000-1.212) |
| **48hr Baseline** | 753 | 57394 | 58147 | .0131 |  |
| **DMRs hypomethylated** |  |  |  |  |  |
| **24hr BFT2** | 993 | 68422 | 69415 | .0145 | 1.112 (1.006-1.230) |
| **24hr Baseline** | 631 | 48351 | 48982 | .0131 |  |
| **48hr BFT2** | 757 | 66188 | 66945 | .0114 | 1.081 (0.971-1.203) |
| **48hr Baseline** | 609 | 57538 | 58147 | .0106 |  |

<sup>a</sup> Odds represent the number of overlapping peaks divided by the number of non-overlapping peaks
<sup>b</sup> Odds ratios represent the odds in BFT2-treated cells divided by the odds in untreated cells
<sup>c</sup> 95% confidence intervals were calculated using a chi-square test
Common DMRs and SNVs in CRC were extracted from the COSMIC database

In contrast, our DMR analysis found that a larger proportion of BFT2-induced chromatin accessibility peaks overlap with common DMRs in CRC 48 hours [OR(95%CI): 1.094(1.018-1.176)], but not 24 hours [OR(95% CI): 1.037(0.965-1.114)] after BFT2 treatment (Table 4). This result supports a commonly held paradigm that chromatin changes come first, followed by methylation changes (22). When we further parsed the data to analyze regions of hypermethylation and hypomethylation individually, the results differed slightly, but consistently showed a larger proportion of peaks overlapping with common DMRs in CRC after BFT2 treatment (Table 4). Thus, analysis of DMRS supported our hypothesis that BFT2-induced regions of increased chromatin accessibility are *more* likely to contain sites of differential methylation found in CRC.

## Discussion

Our data support that acute BFT2 treatment of colon epithelial cells (CECs) leads to rapid onset and dynamic, but limited changes in chromatin accessibility. These changes in chromatin accessibility could be associated with BFT2-induced changes in gene expression. We observed changes in both chromatin accessibility and gene expression over a 48-hour period, with the strongest changes occurring at 24 hours, then relaxing to baseline by 48 hours. Our results corroborate previous observations about the acute onset of BFT2 action on epithelial cells in vitro and in vivo (7, 16, 18).

Our RNA-seq data identifies several genes and pathways that are differentially regulated after BFT2 treatment. The pathways identified, particularly at 48 hours, include those involved in the DNA damage response and cell cycle regulation, both germane to previously reported BFT-induced CEC changes in vitro and in vivo. Namely, BFT2 has been shown to induce DNA double strand breaks and modulate apoptosis *in vitro* (17, 20) and ETBF similarly induces DNA double strand breaks (16) as well as colon carcinogenesis *in vivo* (23). In addition, our study also identified several new genes that may explain how BFT2 contributes to tumorigenesis by allowing other bacteria to invade the mucus layer and bind to CECs. Specifically, BFT2 causes upregulation of CEACAM6 and MUC16, and downregulation of MUC2. CEACAM6 acts as a receptor for adherent-invasive *E.coli*, a particular strain of *E. coli* that has recently been implicated in the development/progression of CRC in individuals with FAP (24) and has previously been associated with sporadic CRC (25). MUC2, the major MUC protein present in human colonic mucus, is notably downregulated after BFT2 treatment, potentially contributing to mucus invasion by tumorigenic *E.coli.* The upregulation of MUC16, a particular MUC variant that is not normally expressed in the colon, may represent a compensatory mechanism (i.e., an attempt to protect CECs after MUC2 downregulation). Collectively, these results may begin to explain how ETBF works with other bacteria to promote tumor formation and growth.

Several previous studies have shown that chromatin structure in gene promoter regions correlates with gene expression (26, 27). Our study took this a step further by showing that acute changes in chromatin accessibility correlate with acute changes in gene expression. 24 hours after BFT2 treatment, increases in chromatin accessibility generally correlated with increases in gene expression while decreases in chromatin accessibility correlated with decreases in gene expression. On the other hand, at 48 hours, no significant association is seen between changes in chromatin accessibility and gene expression. This lack of association is likely due to the limited gene expression changes that remain 48 hours after BFT2 treatment and underscores a trend that is seen throughout most of our data whereby changes due to BFT2 treatment are greater 24 hours after treatment and approach baseline levels by 48 hours. This pattern is also corroborated by previous *in vitro* data that has shown that the effect of BFT2 on gene expression in CECs is rapid and appears to decay quickly (7, 16, 18), although biologic effects such as CEC proliferation persist for at least 72 hours *in vitro* (7).

Areas of increased chromatin accessibility after BFT2 treatment are enriched in binding motifs for many transcription factors belonging to the AP-1/ATF family of transcription factors, including FOSL1, JUN and JDP. These transcription factors function downstream of the JNK pathway known to be activated by BFT2 (18, 20, 21) . After BFT2 treatment, we expected to see increased expression of genes which are regulated by these particular transcription factors, but chromatin opening alone appeared insufficient to drive changes in gene expression of these JNK pathway genes. Hence, we looked for transcription factors that had both an enriched binding motif and increased gene expression after BFT2 treatment. FOSL1 was the primary candidate revealed by our analysis. We found that though presence of a FOSL1 binding site alone was unable to predict differential gene expression after BFT2 treatment, when coupled to promoter chromatin accessibility, an excellent concordance with increased gene expression was detected. This finding exemplifies how integration of chromatin accessibility and gene expression data can be applied to understand how BFT2 acts on CECs. We needed to use both types of data to correctly surmise that transcription factor upregulation, enrichment of transcription factor binding motifs, and the presence of open chromatin in the promoter region all contribute to BFT2-induced gene regulation.

Because CRC usually results from a series of genetic mutations, and can be transformed by changes in methylation patterns, any potential interaction between BFT2, chromatin accessibility, and common SNVs/DMRs in CRC is important. Herein, we found a BFT2-induced *increase* in chromatin accessibility at sites of common DMRs in CRC and a *decrease* at sites of common SNVs in CRC. Increased methylation after inoculation of mice with ETBF has been suggested previously. Specifically, studies by O’Hagan and colleagues have shown that binding of DNMT1 (DNA methyltransferase 1) at promoters of low-expressing genes is upregulated after ETBF murine colonization (11). While this study implicates ETBF, and thus BFT, in DNA hypermethylation, no studies to date have explored the connection between BFT and hypomethylation. Our data suggest that BFT2-induced changes in chromatin accessibility may have dual effects on chromatin methylation patterns (fig 4). We hypothesize that increased chromatin accessibility is, at least, one step critical to chromatin methylation pattern changes in response to BFT2. This is important because hypermethylation can lead to silencing of tumor suppressor genes and hypomethylation can lead to upregulation of transposable elements that may reintegrate into tumor suppressor or proto-oncogenes. In the future, bisulfite sequencing experiments can be conducted to definitively correlate sites of methylation change after BFT2 treatment with sites of altered chromatin accessibility.

Several experiments, conducted across a variety of species, have also shown that chromatin accessibility may be inversely correlated with rates of specific DNA mutations (28–30). One study in particular showed that DNase accessible euchromatin was protected from UV-induced DNA damage, while lamina-associated heterochromatin was more susceptible to damage (14). This study is of particular importance because it investigated carcinogenesis-related DNA damage. Our data suggest that BFT2-induced decreases in chromatin accessibility are correlated with frequently mutated areas, suggesting that BFT2-induced chromatin changes may increase the chances of mutations. Such a hypothesis would require significant further testing to determine its accuracy – this could be done by sequencing BFT2-treated and untreated cells in order to determine if BFT2 treatment increases the mutation rate. If so, sites of increased mutation frequency could be identified and correlated with sites of BFT2-induced chromatin accessibility change.

Our study has several limitations that provide avenues for future experiments. Most notably, our experiments were performed on HT29/C1 cells, a colon carcinoma cell line. These cells contain several mutations that may lead them to respond differently to BFT2 than normal CECs. As a result, these experiments should be confirmed in a primary cell culture system, or ideally, *in vivo* in a mouse model or from human samples with and without ETBF colonization. Our study, by design, was also performed over only an early time window. *In vitro* studies with BFT best mimic the earliest *in vivo* events whereby ETBF induces CEC changes and colitis by ~24 hours after murine colonization (31). In contrast, the persistent CEC exposure to BFT afforded by chronic ETBF colonization *in vivo* cannot be reliably modeled *in vitro*, in part, because of CEC uptake and degradation of BFT (32). Thus, additional *in vivo* studies are warranted since ETBF murine colonization is persistent and associated with ongoing IL-17-dominant inflammation and CEC hyperplasia after one year in C57Bl/6 mice (31, 33). Ultimately, we want to know which BFT-induced chromatin and gene expression changes persist and contribute to tumor formation. Verifying these experiments in an in vivo system will help answer some of the questions proffered by these data.

The experiments reported herein provide additional insight into the effect of BFT2, a bacterial exotoxin linked to CRC pathogenesis, on CECs. We have identified new associations between BFT2-induced chromatin accessibility and gene expression changes, and also correlated these changes with previously published data on BFT2-induced signal transduction and DNA modifications that may contribute to tumor formation or growth. We need to better understand how BFT2 affects the genome and epigenome of CECs to determine if asymptomatic carriers of ETBF are at increased risk for colon tumorigenesis.

## Acknowledgements

This work was supported by the Bloomberg Philanthropies (CLS), National Institutes of Health R01CA179440 (CLS), the Johns Hopkins Department of Medicine (CLS), and Howard Hughes Medical Institute (JA)

All authors declare no completing interests.

## Materials and Methods

### Culture of HT-29/C1 cells

HT29/C1 cells were cultured at 37°C and 10% CO2. Cells were plated at 20% confluency 4 days before addition of BFT2 toxin. At day 0, HT29/C1 cells were washed five times with PBS, placed in minimal media (DMEM) without FBS or Pen/Strep and toxin was added one time at a concentration of 100ng/ml. After toxin addition, cells were allowed to grow at 37°C and 10% CO2 for 24 or 48 hours.

### ATAC-seq transposition reaction

For all experiments, cell counts were obtained by trypsinizing cells in 0.5% trypsin for 10 minutes at 37°C and then counting using a hemocytometer. Non-trypsinized cells were then scraped and 50,000 cells were added to an Eppendorf tube. ATAC-seq was performed using the protocol outlined by Buenrostro et al. (2015). Briefly, cells were washed with 50uL cold 1x PBS buffer, then centrifuged at 500g for 5 minutes at 4°C. Cells were then resuspended in 50uL cold lysis buffer (10 mM Tris-HCl [pH 7.4], 10 mM NaCl, 3 mM MgCl_2_, 0.1% IGEPAL CA-630), gently lysed to preserve cell nuclei, and centrifuged again at 500g for 10 minutes at 4°C. Cells were then washed 3x with wash buffer (lysis buffer without igepal). After each wash, cells were centrifuged at 500g for 5 minutes at 4°C. Cell nuclei were then resuspended in the transposition reaction mix and incubated for 30 minutes at 37°C. DNA was purified using a Qiagen MinElute PCR Purification Kit and eluted in 10uL elution buffer. DNA was stored at −20°C until fragments were amplified via PCR.

### PCR amplification following transposition

All of the DNA purified following transposition was PCR amplified. To do so, the following were combined in a 0.2mL PCR tube: 10uL transposed DNA, 10uL nuclease-free H2O, 2.5uL 25uM custom Nextera PCR primer 1, 2.5uL 25uM custom Nextera barcoded PCR primer 2, 25uL NEBNext high-fidelity 2x PCR master mix. The components were amplified as follows: 1 cycle of 72°C for 5 min, 98°C for 30 sec; 5 cycles of 98°C for 10 sec, 63°C for 30 sec, 72°C for 1 min. After initial amplification, the number of additional cycles to run was determined using qPCR. For this, the following were combined in a 0.2mL PCR tube: 5uL of previously PCR-amplified DNA, 4.5uL of nuclease-free H2O, 0.25uL of 25uM custom Nextera PCR primer 1, 0.25uL of 25uM custom Nextera PCR primer 2, 5uL KAPA SYBR FAST qPCR master mix (2x) (total reaction 15 ul). The components were amplified as follows: 1 cycle of 98°C for 30 sec; 20 cycles of 98°C for 10 sec, 63°C for 30 sec, 72°C for 1 min. To calculate the number of additional cycles of PCR needed, linear Rn versus cycle was plotted and the cycle number that corresponds with 1/3 of the maximum fluorescent intensity was determined. After the number of additional PCR cycles was determined, the remaining 45uL of PCR product was run as follows: 1 cycle of 98°C for 30 sec; N cycles (as determined via qPCR) of 98°C for 10 sec 63°C for 30 sec, 72°C for 1 min. Finally, the amplified library was purified using 0.9x Ampure beads at room temperature, and eluted in 20uL RNAse-free H2O.

### ATAC-seq data analysis

The 4nM pooled ATAC-seq library was sequenced on an Illumina HiSeq using 50bp paired-end sequencing. For each sample, three biological replicates were sequenced. After sequencing, the data was analyzed using the pipeline developed by the Kundaje lab, as outlined on ENCODE (35). Briefly, reads were trimmed, aligned and filtered using Bowtie2(36) and peaks were called using MACS2(37). The three replicate peak files were combined in order to create one consensus file for each treatment condition using an irreproducible discovery rate (IDR) threshold of 0.1. For all analyses, the “optimal set” consensus file was used. Peak files for samples before and after BFT2 treatment were then compared in order to create BFT2-opened peaks (peaks present after BFT2 treatment, but not before) and BFT2-closed peak (peaks present before BFT2 treatment, but not after) files at 24 and 48 hours after treatment. The ChIPpeakAnno package was used to separate peaks into specific regions of the genome (promoter, coding, intergenic) (38). The ChIPpeakAnno package was also used to associate peaks located in promoter regions with their nearest downstream genes. The haystack_bio package was used to identify transcription factor binding motifs (39). For this analysis, C-G correction was turned off. For both the 24 hours and 48 hours analyses, the BFT2-treated peak file was compared to the baseline peak file. A ratio greater than 1 represents transcription factor binding motifs that are present more frequently in the BFT2-treated peak file, while a ratio of less than 1 represents transcription factor binding motifs that are present more frequently in the baseline peak file. The bedR package was used to calculate the jaccard index for the overlap between identified peaks in our dataset and CTCF binding peak data in Caco-2 cells, taken from the UCSC genome browser (40).

### RNA-seq assay

Cells were washed once with PBS, collected using a cell scraper, and RNA was extracted from cells using the Qiagen RNeasy mini kit. After RNA extraction, the RNA pellet was flash frozen using 100% ethanol and dry ice. Samples were then stored at −80^<u>o</u>^C. For library preparation, samples were removed from storage, and mRNA was enriched using the NEBNext Poly(A) mRNA Magnetic Isolation Module. Afterwards, a non-directional RNA-seq library was constructed using the NEBnext Ultra RNA Library Prep kit from Illumina. The 2nM pooled RNA-seq library was sequenced using the Illumina HiSeq. For one sample, 50bp paired-end sequencing was performed. For the other two samples, 50bp single-end sequencing was performed.

### RNA-seq data analysis

After sequencing, Kallisto was used to perform pseudoalignment of the raw RNA-seq data (41). Then, Sleuth was used to quantify gene expression and perform differential expression analyses (42).

### Gene Set Enrichment Analysis

Gene set enrichment analysis was performed using all genes with a P-value < 0.05 at 24 or 48 hours after BFT2 treatment. For this analysis, a ranklist was first created by sorting the differentially expressed genes using the following formula: sign of fold change × inverse P-value. This created a rank ordered list with the most significantly upregulated genes at the top and the most significantly downregulated genes at the bottom. The GSEA Pre-ranked tool was then run, using 1000 permutations, a classic (instead of weighted) analysis, a minimum gene set size of 15 and a maximum gene set size of 1500. The output of this analysis was then converted into graphical and tabular formats.

### Statistics

To calculate the correlation between chromatin accessibility (ATAC-seq) and gene expression (RNA-seq) at baseline and after BFT2 treatment, the Wilcoxon Rank Sum test was used with an alpha of 0.05. To calculate the enrichment of transcription factor binding motifs in BFT2-treated and untreated samples, a Fisher’s exact test with an alpha of 0.01 was used. To determine if there was a correlation between the location of BFT2-treated peaks and common SNVs and DMRs in CRC, a chi-square test with an alpha of 0.05 was used in which BFT2-treated peak files were compared to baseline peak files.

### Accession number(s)

The complete experimental data set was deposited in the Gene Expression Omnibus (GEO) database under accession number GSE113220.

